# Autophagy coordinates chondrocyte development and early joint formation in zebrafish

**DOI:** 10.1101/2021.08.13.456237

**Authors:** Joanna J. Moss, Martina Wirth, Sharon A. Tooze, Jon D. Lane, Chrissy L. Hammond

## Abstract

Autophagy is a catabolic process responsible for the removal of waste and damaged cellular components by lysosomal degradation. It plays a key role in fundamental cell processes, including ER stress mitigation, control of cell metabolism, and cell differentiation and proliferation, all of which are essential for cartilage cell (chondrocyte) development and survival, and for the formation of cartilage. Correspondingly, autophagy dysregulation has been implicated in several skeletal disorders such as osteoarthritis and osteoporosis. To test the requirement for autophagy during skeletal development in zebrafish, we generated an *atg13* CRISPR knockout zebrafish line. This line showed a complete loss of *atg13* expression, and restricted autophagic activity *in vivo*. In the absence of autophagy, chondrocyte maturation was accelerated, with chondrocytes exhibiting signs of premature hypertrophy. Focussing on the jaw element, autophagy disruption affected joint articulation causing restricted mouth opening. This gross behavioural phenotype corresponded with a failure to thrive, and death in homozygote *atg13* nulls within 17 days. Taken together, our results are consistent with autophagy contributing to the timely regulation of chondrocyte maturation and for extracellular matrix formation.

## INTRODUCTION

Macroautophagy (henceforth termed autophagy) is a catabolic process which enables the breakdown of cytosolic components—including misfolded protein aggregates, redundant organelles, and invading microorganisms—into their basic biomolecular constituents by the actions of lysosomal acid hydrolases. This cellular recycling pathway is essential during differentiation, and contributes to cell and tissue homeostasis, where its primary function is to mobilise nutrients to sustain vital cellular functions during stress ^1^. It involves the formation of an isolation membrane that sequesters cytoplasmic cargo, and this membrane structure finally seals to form a new organelle—the autophagosome—which is subsequently delivered to the lysosome for degradation. This process is mediated by a collection of autophagy-related (ATG) proteins which can be categorised into four functional complexes: the ULK (Unc-51-like autophagy-activating kinase) complex; the phosphatidyl inositol 3-kinase complex I (PI3KC3); and the two conjugation systems, ATG5–ATG12-ATG16L and MAP1LC3–ATG3 ^2, 3^.

In vertebrates, autophagy initiation requires the formation of the ULK complex, comprising the serine/threonine protein kinase ULK1/2, and its adaptors, ATG13, FIP200, and ATG101 ^4^. Upon autophagy induction, the ULK complex translocates to discrete foci dispersed throughout the cell, typically associated with endoplasmic reticulum (ER) membrane, where it phosphorylates components of PI3KC3, triggering recruitment of the remaining core autophagy machinery ^2, 5^. Along with FIP200, ATG13 has been shown to play a vital role in both the localised activation of the ULK1 kinase and its recruitment at the nascent isolation membrane site ^6, 7^. Here, ATG13 helps form a building scaffold for other proteins within the autophagy pathway to bind and be stabilised ^8^. These roles require the two key domains that comprise ATG13: the C-terminal HORMA (Hop1/Rev7/Mad2) domain, and the N-terminal IDR (intrinsically disordered region) ^9, 10^. In particular, the highly conserved HORMA domain is essential for autophagy induction and the recruitment of PI3KC3 via ATG14 ^11^. In a range of vertebrate models and cell lines, loss of ATG13 expression has been shown to block autophagy activity (^12–17^; see table 3 in ^18^). Meanwhile, mouse knockout models for Atg13 are embryonic lethal, a result consistent with the targeting of other autophagy genes in murine knockout models, including Fip200, and Atg7 ^19–21^. Interestingly, mouse knockout models have demonstrated that, generally, loss of upstream autophagy genes causes lethality earlier in development compared to *atg* genes encoding members of the conjugation systems: Beclin1 and Vps34 (in addition to those mentioned above) are embryonically lethal ^22, 23^, whereas genes acting later (Atg3, Atg5, Atg7, Atg9a, Atg12, Atg16L1 and by contrast, Ulk1/2) show neonatal lethality (see Table 1 in ^24^). A result also mirrored in knockout zebrafish lines for the equivalent *atg* genes ^25^. Therefore, these results demonstrate that autophagy and its machinery are involved in multiple cellular processes beyond metabolic recycling, and are thus essential for survival ^26^.

Extensive studies have established the importance of autophagy in a range of housekeeping pathways, such as the control of cell metabolism, mitigation against endoplasmic reticulum (ER) and mitochondrial stress, in addition to roles during the coordination of cellular differentiation and proliferation ^27–29^. Each of these processes have been shown to be essential for cartilage cell (chondrocyte) development and survival, and for chondrogenesis – the formation of cartilage from condensed mesenchymal tissue ^30–32^. Correspondingly, autophagy dysregulation has been implicated in several skeletal disorders, including the degenerative joint disorder, osteoarthritis ^32–36^. Osteoarthritis (OA) is the most common cause of arthritis in the world ^37, 38^ and one the biggest causes of disability, causing an increasing global economic burden through lost working days and forced early retirement ^39^. OA affects all structures of the joint and is characterised by the progressive degeneration of cartilage causing, synovial inflammation, osteophyte formation, ligament damage, bone misalignment and joint pain ^40, 41^. Although often characterised as a disease of ageing, recent studies have highlighted the effect of improper joint shape formation during skeletogenesis on OA development in later life ^42–45^. Therefore, determining the impact of cellular processes, such as autophagy, on the co-ordination of cartilage and joint development is essential for deepening our understanding of the pathogenesis of OA and how this disease can be best treated.

The process of chondrogenesis begins with the condensation and differentiation of mesenchymal stem cells at sites of future skeletal formation. These cells differentiate into cartilage forming chondrocytes which secrete a characteristic extracellular matrix (ECM) formed largely of type II collagen α1 (Col2a1) and specific proteoglycans, such as aggrecan, generating a cartilage matrix ^46^. As chondrocytes develop, they pass through a well characterised set of maturation steps which include intercalation and proliferation, where cells flatten and separate into a narrow, single-cell stacked column, followed by hypertrophication, as cells exit the cell cycle and switch from secreting Col2a1 to type X collagen α1 (Col10a1) ^47–49^.

Several studies have identified important roles for autophagy throughout chondrogenesis. In the early stages of this process, *in vitro* studies have shown a positive correlation between autophagy activity and chondrocyte proliferation and differentiation ^50^, whilst maturing chondrocytes show high MAP1LC3 expression ^51^. In mice, during early chondrogenesis, chondrocyte autophagy is induced in the growth plate postnatally ^52^, and the conditional loss of Atg5 or Atg7 during chondrogenesis has been shown to reduce growth plate activity and cause growth retardation ^50^, which is likely triggered by ER stress within chondrocytes ^53^. Additionally, conditional knockout of Atg7 in mouse chondrocytes causes enlarged ER cisternae, extracellular matrix (ECM) disorganisation and retention of procollagen 2 (a preform of Col2a1 – a major component of cartilage matrix), as well as reduced chondrocyte proliferation and survival ^52, 53^. If autophagy is disrupted later in chondrogenesis, chondrocytes in the proliferative stage show an accumulation of glycogen granules, severe growth retardation, and increased apoptosis ^30^. Meanwhile, looking beyond early development, reduced autophagy activity via a chondrocyte-specific Atg5 deletion causes the premature onset of osteoarthritis development in mice from 6 months ^34^. Together these studies demonstrate a key role for autophagy in chondrocyte proliferation and cartilage growth; and highlight how disruption to this process can lead to the premature development of degenerative joint disease. However, the role of the autophagy pathway within chondrocyte differentiation is still incompletely understood.

In this study, we have generated a new *atg13* knockout zebrafish line and characterised its role during cartilage and joint development. We find that suppression of autophagy accelerates chondrocyte maturation, leading to improper chondrocyte intercalation and disruption to jaw joint formation and movement. We argue that this developmental defect contributes to lethality at around day 17, due to a failure of *atg13* knockout fish to thrive at free-feeding stages. The disruption to joint function seen in this model indicates an important role for autophagy in supporting the regulation of chondrocyte development to ensure proper joint formation and highlights an important pathway through which autophagy dysregulation may contribute to OA development.

## METHODS

### Zebrafish Husbandry and Transgenic Lines

Zebrafish were raised and maintained under standard conditions ^54^. Experiments were performed under UK Home Office project licences, under approval by the local ethics committees (the Animal Welfare and Ethical Review Body of the University of Bristol and the Francis Crick Institute). Transgenic lines *Tg(CMV:EGFP-map1lc3b)* ^55^ and *Tg(Col2a1aBAC:mcherry)* ^56^ have been reported previously.

The *atg13* stable zebrafish line (*atg13^fci500^*) was generated using CRISPR-Cas9 mutagenesis. Briefly, gRNAs targeting exon 3 of the zebrafish *atg13* orthologue (Ensembl: ENSDART00000052324.6; zgc:63526) were cloned into the pT7-gRNA plasmid (Addgene #46759) and generated according to Jao *et al*. ^57^; CRISPR1-atg13 (exon 3)-For: 5’-<u>TAGG</u>TTATAGTGCAAGCCCGGCT-3’ and CRISPR1-atg13 (exon 3)-Rev: 5’-<u>AAAC</u>AGCCGGGCTTGCACTATAA-3’. Following generation of Cas9 mRNA using the pT3TS-nls-zCas9-nls plasmid (Addgene #46757) ^57^, Cas9-encoding mRNA (200 ng/μl) and *atg13* targeting gRNA (~16-25 ng/μl) were co-injected into one-cell stage embryos. High-resolution melting analysis (HRMA) ^58^ was performed on DNA from single embryos and fin clips to determine the efficacy of mutagenesis in F0 and F1 embryos and to identify adult F1 heterozygous carriers. The PCR product sequenced and a 5-base pair (bp) deletion at 92-96 bp (31aa, zv9 chr19:7834334) in the *atg13* gene was detected, causing a premature stop codon and insertion of a novel HindIII restriction site at the mutation site.

For autophagy flux analysis, larvae were treated at 4dpf or 5dpf with 100 nM BafilomycinA1 (BafA1; 14005, Cambridge Bioscience, UK) or DMSO diluted in Danieaus for 3 hours at 28°C.

### LysoTracker live staining

For visualisation of lysosomal compartments, LysoTracker Red DND-99 (L-7528, Invitrogen, MA, USA) was used. Following treatment with DMSO or BafA1, *Tg(atg13;CMV:EGFP-map1lc3b)* 4dpf larvae were placed in 10 μM LysoTracker Red in Danieaus for 1 hour at 28°C and then washed three times (3 minutes per wash) in fresh Danieaus. LysoTracker-positive and GFP-Lc3 puncta were counted from single z-slices from three independent fields of a set size per larvae.

### Cell proliferation assay

Proliferation of the larvae was measured using the Click-iT EdU imaging kit (Invitrogen) according to the manufacturer’s instructions. Briefly, *Tg(atg13;Col2a1aBAC:mcherry)* larvae at 5dpf were treated with 400 μM EdU in Danieaus for 24 hrs. Larvae were fixed in 4% PFA and then incubated in the Click-iT reaction cocktail for 30 min.

### DNA extraction and genotyping

Whole embryos or caudal fin amputations on larvae ^59^ at 3-5dpf were incubated in base solution (25 mM NaOH, 0.2 mM EDTA) for 30 minutes at 95°C before addition of equal volume of neutralization solution (40 mM Tris–HCl, pH 5.0). For genotyping, touchdown PCR (with primers *atg13* F- GGCTCGTGCGACAATGGATAGTG; R- GACCTCGGGGATGTCCTTTATTGC) was followed by a HindIII restriction digest (R3104S, New England Biolabs, MA, USA), and fragments were separated by gel electrophoresis on a 3% agarose gel. Digestion product for wildtype PCR product is 386 bp; 201 bp for *atg13* mutant allele.

### Western blotting

Following treatment with DMSO or BafA1 at 5dpf, larvae were deyolked and snap frozen in liquid nitrogen. Hot 3x SDS sample buffer (100 μl per 15 larvae at 5dpf) was added and samples homogenised using by pulling sample through a 0.22 gauge needle on ice 6-8 times. Lysates were heated at 95°C for 10 min and resolved on 10% or 12% polyacrylamide gels at 40mA after loading 25 μl per sample. Following transfer at 100V, nitrocellulose membranes were blocked in 5% milk-TBT-1% Triton, then incubated with primary antibodies diluted in 5% milk overnight at 4°C (anti-ATG13 (SAB4200135, 1:100, Sigma, MI, USA); anti-GAPDH (10494-1-AP, 1:2000, Proteintech, IL, USA); anti-LC3B (ab51520, 1:300, Abcam, Cambridge, MA, USA); anti-p62/SQSTM1 (5114, 1:300, Cell Signalling Technologies, MA, USA)). After washing in TBS-T, blots were incubated with secondary anti-rabbit (G33-62G, 1:10,000, SignalChem, Canada) and anti-mouse (G32-62G, 1:10,000, SignalChem) horseradish peroxidase-conjugated antibodies at room temperature for 1 hr before exposure on photographic film (28906844, Amersham, UK) following incubation with ECL for 1 minute (RPN2232, GE Healthcare, IL, USA) as per manufacturer’s instructions.

### Whole-mount Immunohistochemistry

Larvae were fixed in 4% PFA then stored at −20°C in 100% methanol. Larvae were re-hydrated, washed in PBS-T before permeabilization with proteinase K (4333793, Sigma) at 37°C. For A4.1025 antibody staining, larvae were instead permeabilised in 0.25% Trypsin for 15 minutes on ice. Larvae were then blocked in 5% horse serum and incubated with primary antibodies overnight at 4°C, unless otherwise stated (anti-myosin (clone A4.1025) (sc-53088, 1:500, Insight Biotech, UK); anti-GFP (ab13970, 1:200, Abcam); anti-mCherry (M11217, 1:100, Invitrogen); anti-col2a1 (II-II6B3-s, 1:20, DSHB, IA, USA); anti-colXa1 (SAB4200800, 1:100, 48 hr incubation needed, Sigma); anti-sox9a (GTX128370, 1:300, Genetex, CA, USA)). Larvae were washed extensively before incubation with Alexa-Fluor secondary antibodies (Invitrogen) diluted in 5% horse serum at 1:500 for 2 hrs at room temperature in the dark. For DAPI staining, larvae incubated in PBS with 1% Triton (PBS-T) containing DAPI (1 μg/mL) for 1 hour at room temperature before washing.

### Confocal Microscope Imaging of Larvae

For confocal imaging, larvae were mounted ventrally in 1% LMP agarose (16520050, Thermofisher, MA, USA) and imaged using a Leica SP5-II AOBS tandem scanner confocal microscope attached to a Leica DMI 6000 inverted epifluorescence microscope and oil immersion 20X or 40X objectives. The microscope was located in the Wolfson Bioimaging Facility, Bristol and run using Leica LAS AF software (Leica, Germany). Maximum projection images were assembled using LAS AF Lite software (Leica) and Fiji ^60^.

### Stereomicroscope Imaging of Zebrafish

Images of live larvae from 1-7dpf were obtained using a Leica MZ10 F modular stereo microscope system at 1-8.3x magnification. For live imaging, fish were anaesthetised using 0.1 mg/ml MS222 (Tricaine methanesulfonate) diluted in Danieaus and imaged laterally.

### Measurement of Jaw Movement Frequency

High-speed movies of 1000 frames were made of jaw movements of *wt* and *atg13* mutants at 5dpf. The number of mouth movements was recorded per 1000 frames for 5 *wt* and *atg13* mutants, respectively. Frames displaying the three widest jaw openings were selected per fish and the distance between the lower and upper jaw at each joint calculated in μm.

### Transmission electron microscopy (TEM)

Following treatment with DMSO or BafA1 for 3 hours, larvae at 5dpf were fixed in 2.5% glutaraldehyde in 0.1 M sodium cacodylate buffer (pH 7.3) at 4°C, washed and then fixed in 0.2 M osmium in sodium cacodylate buffer with 1.5% ferrocyanide for 1 hr at room temperature. After washing, larvae were placed into sample processor for transmission electron microscopy (TEM) using a standard Epon resin protocol. Briefly, larvae were postfixed in reduced osmium, stained with uranyl acetate dehydrated in ethanol and infiltrated with Epon resin via propylene oxide, and polymerised at 60°C for 48 hours.

Ultra-thin sections (50 nm) of epon embedded larvae were cut using a diamond knife, collected on Formvar coated one-hole copper grids (AGS162, Agar Scientific, UK) and observed using a Tecnai 12-FEI 120kV BioTwin Spirit transmission electron microscope. Images of chondrocytes from transverse sections of the ethmoid plate were collected, (n= 3 larvae per genotype, per condition) using an FEI Eagle 4k × 4k CCD camera and analysed using the freehand selection tool and multi-point counter in Fiji ^60^.

### Image analysis and statistics

Larval lengths were obtained from lateral images at 1-7 dpf by measuring from the tip of the mouth to the end of the tail manually in ImageJ.

Whole jaw measurements and muscle fibre number and length measurements were taken manually in ImageJ from max projections of confocal z-stacks of *wt* and *atg13* mutant larvae immunostained with Col2a1 or A4.1025 antibodies, respectively.

Modular cell analysis was performed using the freely available Modular Image Analysis (MIA; version 0.9.30) workflow automation plugin for Fiji developed by Dr. Stephen Cross ^60, 61^. Program requires input of confocal z-stacks of larval jaw joints, labelled for Col2a1 to allow for the calculation of cell number and volume, jaw element volume and distance, and inter-element volume.

To quantify Sox9a expression, the Seg3D program was used (a custom script written in MATLAB (version 2015a; Mathworks) developed by Dr. Stephen Cross and previously described ^62^) whereby the volume of Sox9a positive expression within the Col2a1 positive cells is calculated using confocal z-stacks in specified regions of interest which are selected via a freehand tool within the program. Same threshold value was used each individual sample and calculated by averaging the mean of the automatic threshold value given for each stack for *wt* and *atg13* mutants, respectively.

Statistical analyses were performed using Graphpad Prism v.9. Error bars on all graphs represent the mean ± standard deviation.

## RESULTS

### Establishment and characterisation of the *atg13* knockout zebrafish model

To explore the role of autophagy in bone and joint development, we developed an *atg13* knockout zebrafish line using CRISPR-Cas9 mutagenesis. The line has a 5 bp deletion in exon 3 of *atg13*, resulting in a premature stop codon 97 bp downstream of the mutation site, and the introduction of a diagnostic HindIII restriction site (Figure 1A). The mutation occurs in the highly conserved HORMA domain of Atg13, essential for Atg13 function as a component of the ULK1 complex, and necessary for autophagy initiation ^2^. Accordingly, homozygous a*tg13* mutants show complete loss of Atg13 protein expression, while heterozygotes have no discernible difference in protein expression compared to *wt* (Figure 1B), indicative of increased expression from the single remaining allele. These data confirm that the mutation causes a full knockout of Atg13 in the homozygous mutants. Given this, we used the homozygous *atg13* mutants for the majority of experiments in this study, and henceforth, these will be termed ‘*atg13* mutants.’

**Figure 1.**
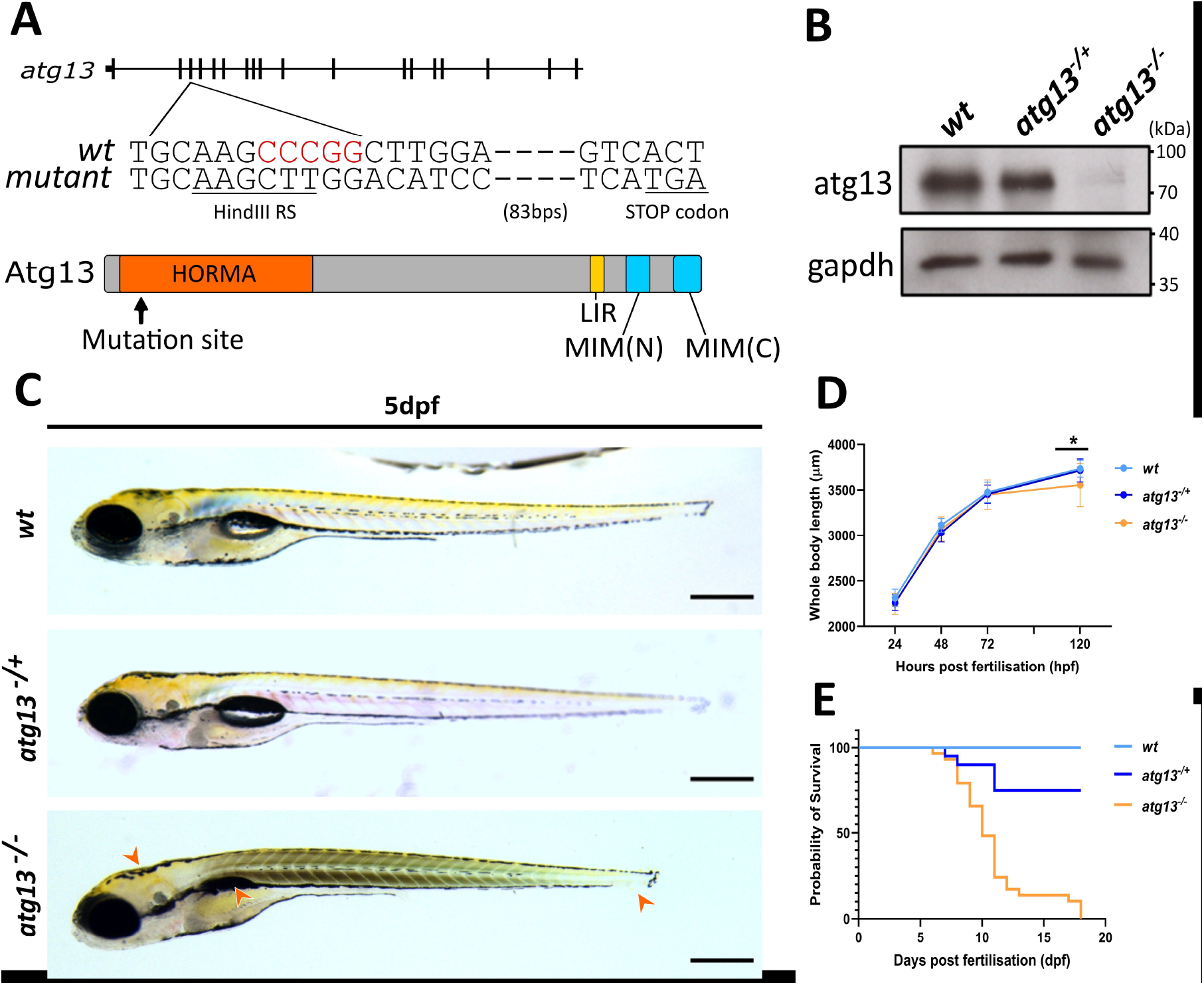
Generation of *atg13* knock out zebrafish line, **(A)** Schematic showing location of 5bp deletion in *atg13* knockout line and generation of premature STOP codon, with black lines representing exons. Key domains within atg13 highlighted; *LIR*, Lc3-interacting region; *MIM*, Microtubule interacting motif. **(B)** Immunoblot showing loss of atg13 expression in *atg13* homozygous mutant. **(C)** Lateral widefield images of *atg13* zebrafish larvae at 5dpf. Orange arrowheads indicate phenotypic differences in development between *wt* and *atg13* mutants (from left to right: bent body axis, uninflated swim bladder and shorter body length). Scale bar = 500μm. **(D)** Graph showing whole body length of *atg13* larvae from 1-5dpf. Student’s t-test performed at 5dpf between *wt* and *atg13* mutant, \**P*=.0013. **(E)** Meire Kaplian graph showing survival of larvae up to 20dpf.

At 24hpf (hours post fertilisation), *atg13* mutant larvae showed no phenotypic differences from *wt*, although by 3dpf certain developmental differences were observed (bent body axis, oedema; supplementary figure S1). By 5dpf, *atg13* mutants were significantly shorter in length and displayed at least one phenotypic difference when compared to *wt* and *atg13* heterozygotes, such as oedema, bent body axis and a failure to fully use yolk sac (indicated by higher yolk sac mass) or inflate the swim bladder, the latter being indicative of a delay in normal development (Figure 1C, D and supplementary figure S1). In line with other mouse and zebrafish knock-out models for key autophagy-related genes ^25, 63^, *atg13* mutants showed juvenile lethality at 17 days post fertilisation (dpf) (Figure 1E). Taken together, these results suggest that Atg13 is essential for zebrafish survival into adulthood, and that loss of Atg13—although not impacting on early survival—disturbs larval developmental rate and morphology.

### *atg13* mutants show reduced autophagy flux

To confirm that autophagy is functionally disrupted in the *atg13* mutants, we generated GFP tagged map1lc3 (GFP-Lc3) zebrafish ^55^ in the *atg13* mutant background (*Tg(cmv:gfp-map1lc3b);atg13)*. Following autophagy initiation, cytosolic Lc3-I is conjugated to phosphatidylethanolamine, a constituent of the phagophore (or isolation membrane), forming lipidated Lc3-II which has greater mobility on SDS-PAGE gels and can be detected in GFP-Lc3 zebrafish as discrete GFP-positive puncta using fluorescence microscopy ^64^. By applying LysoTracker Red, a vital lysosomal dye, we measured the abundance of autophagosomes and lysosomes in *wt* and *atg13* mutants at 4dpf, by counting the number of GFP-Lc3 and LysoTracker positive puncta per cell (Figure 2A). Under basal conditions (DMSO treatment), *atg13* mutants showed no differences in lysosome abundance compared to *wt*, with a non-significant decrease in GFP-Lc3 puncta numbers (Figure 2A and B). Though loss of *atg13* is expected to prevent autophagosome formation through inhibiting early autophagic signalling ^6^, GFP-Lc3 positive puncta were present in *atg13* mutants. GFP-Lc3-positive aggregates have previously been observed in the mouse *atg13* mutant model ^20^ and in *C. elegans atg13/epg1* mutant embryonic cells ^65, 66^ and we propose may also occur in our atg13 mutant zebrafish model.

**Figure 2.**
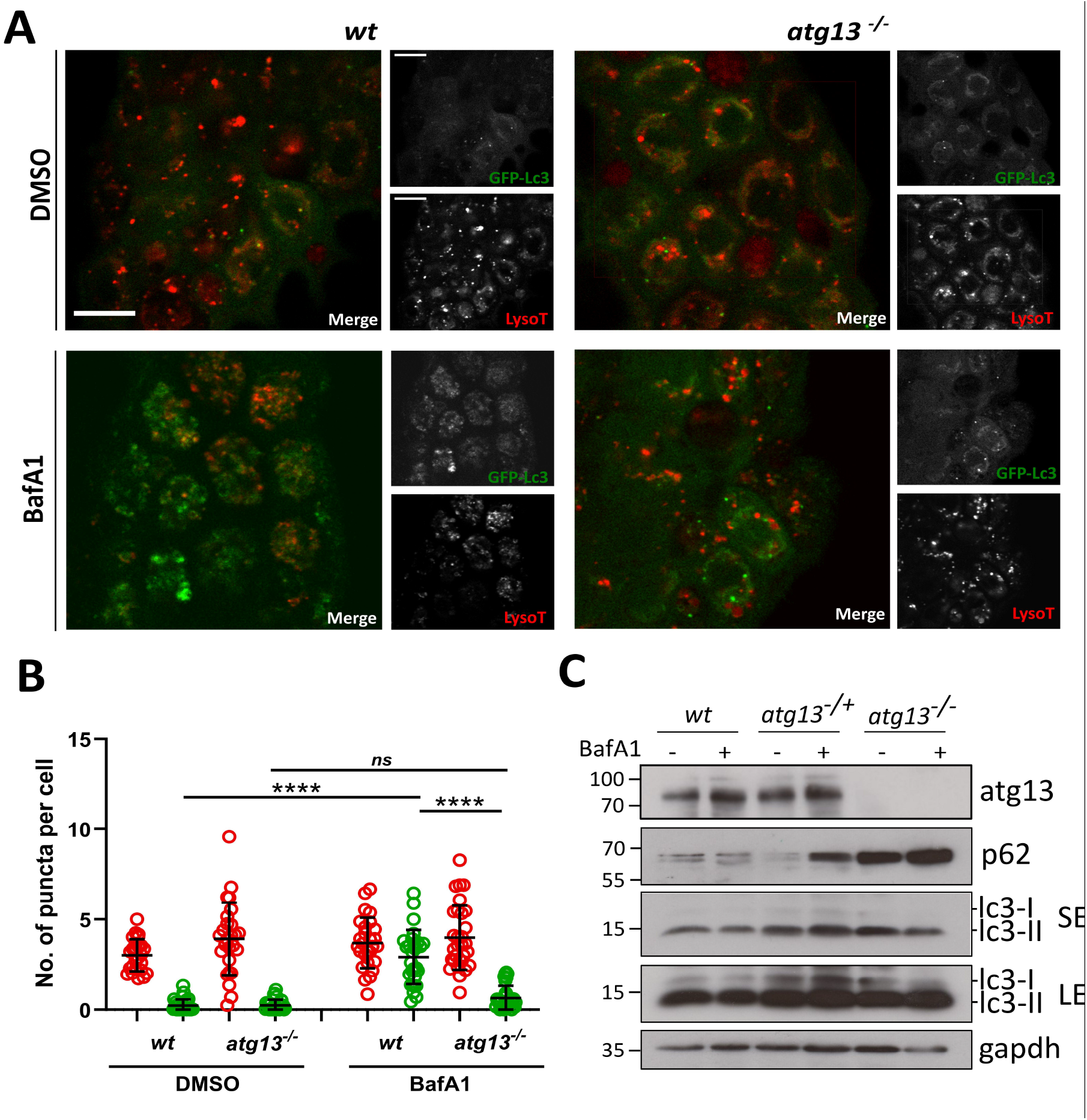
*atg13* mutants show reduced autophagy flux, **(A)** Representative single confocal z-slices of epidermal cells taken from LysoTracker stained *Tg(cmv:gfp-map1lc3;atg13) wt* and mutant larvae, at 4dpf following treatment with DMSO or 100 μM Bafilomycin for 3 hours. Scale bars = 10μm. **(B)** Quantification of number of lysosomal (red) and GFP-Lc3 (green) puncta per cell. 2-way ANOVA performed for each; **** *P*<.0001. **(C)** Representative immunoblot of *atg13 wt*, heterozygous and mutant larvae at 5dpf following treatment with DMSO or 100 μM Bafilomycin for 3 hours. *SE* = short exposure, *LE* = long exposure. Molecular weight markers indicated on right hand side of immunoblots.

Next, to analyse the rate of autophagy flux, larvae were treated with the vacuolar-type H^+^-ATPase inhibitor, Bafilomycin A1 (BafA1). As an inhibitor of lysosomal acidification, BafA1 prevents the formation of autolysosomes, and thus causes accumulation of Lc3-II and autophagic cargo primarily by blocking the degradation of autophagosomes. By comparing Lc3-II puncta numbers and Lc3-II density by immunoblotting under basal and BafA1 treated conditions, autophagic flux rates can be assessed. Following treatment with BafA1, *wt* zebrafish showed a significant increase in GFP-Lc3 puncta numbers compared to basal conditions, indicative of high autophagic flux, as expected (Figure 2A and B). However, there was a dramatically reduced accumulation of GFP-Lc3 puncta in *atg13* mutants, indicating attenuated autophagic flux (Figure 2A and B).

Immunoblot analysis of endogenous LC3 did not reveal clear differences in the abundance of lipidated LC3-II between *wt*, heterozygous and homozygous *atg13* mutants under basal conditions and in response to BafA1 treatment (Figure 2C). In contrast to human or rodent samples, zebrafish appear to show higher basal lipidated LC3-II levels from 2dpf ^55, 67–69^, which could be due to differences in LC3 processing, and we observed a similar pattern even in the absence of atg13 expression. More informative, however, was the analysis of the autophagy receptor protein p62 (SQSTM1), which is degraded by the autophagy machinery, and is thus a good measure of flux. We observed a strong accumulation of p62, indicative of defective autophagic flux in the atg13 mutants (Figure 2C). Taken together, these results demonstrate that loss of *atg13* causes the expected deficiencies in autophagy, correlating with reduced turnover of autophagic substrates. Therefore, under a stress phenotype, *atg13* mutants have a significantly reduced capacity for an autophagy response.

### Joint function is reduced in *atg13* mutant fish

Given the role of autophagy in key skeletal processes, we examined the effect of the *atg13* mutation on cartilage development in the context of joint formation. In zebrafish, skeletal formation begins as early as 2dpf with initial establishment of specific craniofacial cartilaginous structures ^70^. Zebrafish skeletal physiology is comparable to that of mammals, as they share the same joint types and components such as joint cavities, articular cartilage and synovial membranes ^71^. This has been most extensively shown in the larval zebrafish jaw, which has a synovial joint and is often used to model joint development. Therefore, lower jaw elements in zebrafish larvae were used to compare the development of the cartilaginous skeleton template between *wt* and *atg13* mutants.

Zebrafish make two distinct jaw movements for feeding and for breathing, which here we describe as mouth and buccal jaw movements, respectively (Figure 3A; red for mouth movements and yellow for buccal movements) ^72^. For more detailed analysis, videos of jaw movements were taken at 5dpf, and the numbers of movements at either joint were measured alongside the displacement distance between the upper and lower jaw (Figure 3; supplementary videos S2 and S3). We observed that the *atg13* mutants did indeed have a significantly reduced range of motion at both joints compared to *wt*, although the total number of movements remained unaffected (Figure 3B, C). The jaw joint itself showed the greatest reduction in movement in *atg13* mutants indicating that the these have restricted mouth opening which could impede feeding. We performed whole mount immunohistochemistry labelling of myosin in 5dpf larvae to determine whether the different ranges of jaw movements were caused by defects in the jaw muscle architecture (supplementary figure S4A). Crucially, the *atg13* mutants showed no differences in muscle fibre number compared to *wt* (supplementary figure S4B), whilst the width and length of the intermandibularis posterior and interhyal muscles were also comparable (supplementary figure S4C; width data not shown). These data indicate that the changes to jaw movements are not being caused by defects in muscle patterning but are due to altered joint formation and function.

**Figure 3.**
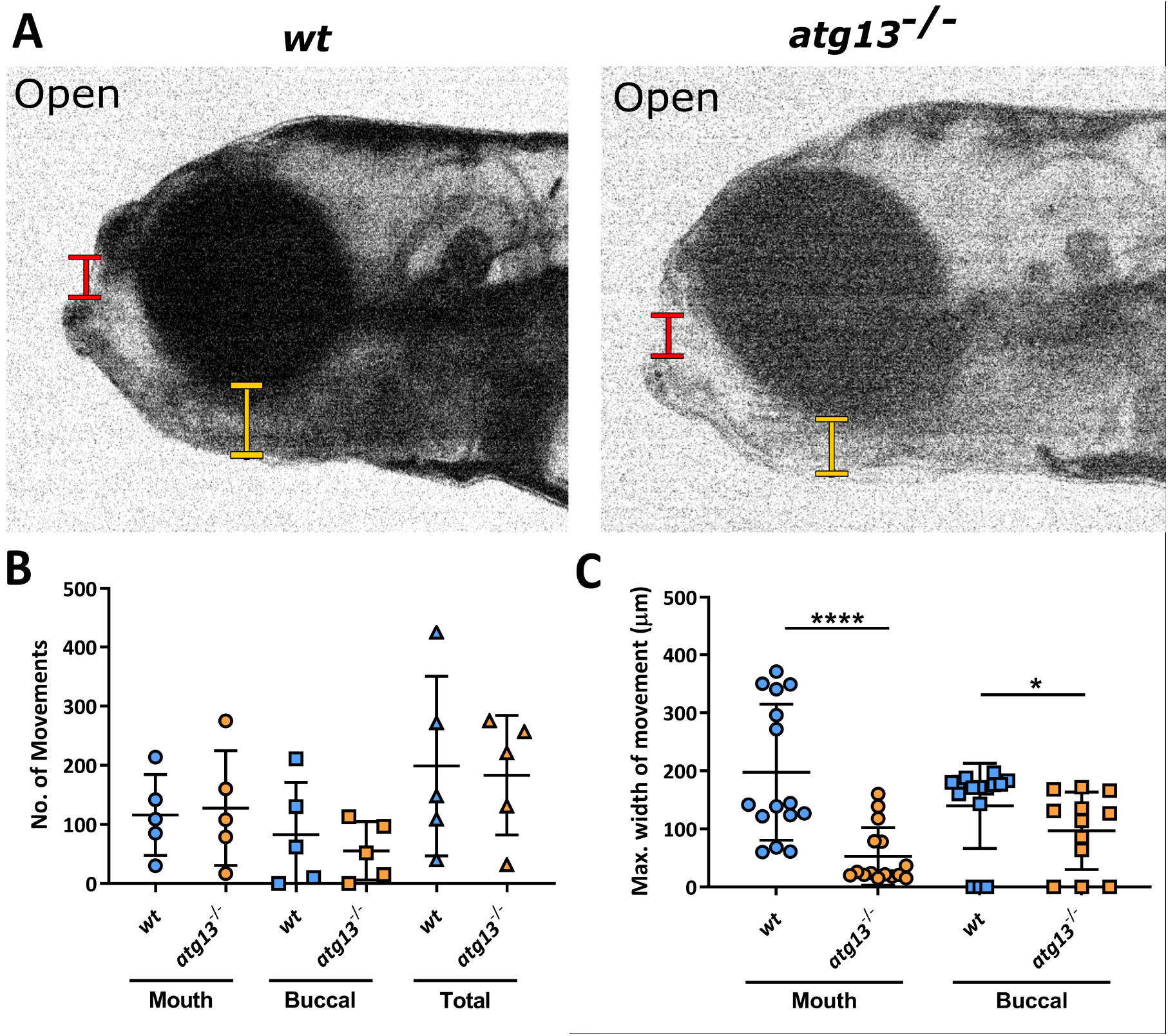
*atg13* mutation reduces jaw function, **(A)** Stills from videos of larval jaw movements taken at 5dpf of *wt* and *atg13* mutant fish. Red and yellow lines indicate where mouth and buccal width measurements taken from, respectively. Quantification of number **(B)** and displacement **(C)** of jaw movements at the mouth and buccal joint. n = 5 for each genotype; three widest jaw openings taken per larvae. Student’s unpaired t test performed for C, **** *P*<.0001, * *P*=.0129.

### Reduced chondrocyte proliferation in *atg13* mutants

Given these changes to jaw opening, we next investigated whether these were due to the loss of autophagy activity affecting the overall growth and formation of the jaw. At 3, 5 and 7dpf, the length and width of lower jaw dimensions were measured using immunostaining of Col2a1 (Figure 4A). from this analysis, it was evident that no changes in Meckel’s cartilage length and width, as well as full length of the lower jaw were observed between the *wt* and *atg13* mutant fish (Figure 4B-D). From this, we inferred that changes to cartilage development caused by the absence of *atg13* appear to be limited to the joint site only and to the cells forming this region.

**Figure 4.**
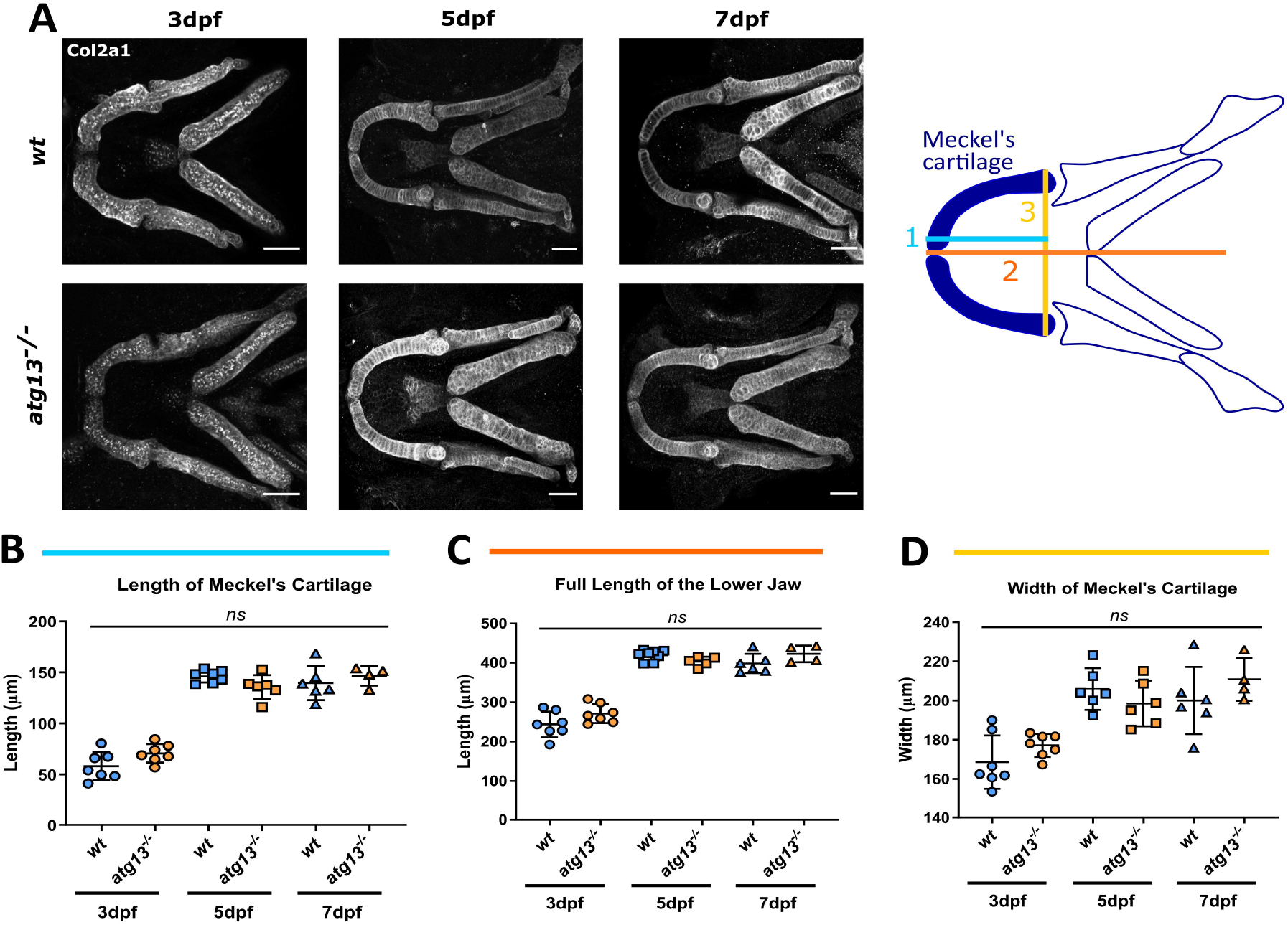
Loss of *atg13* does not affect size of lower jaw in development, **(A)** *Left*, representative confocal z-stack projections of the lower jaw at 3, 5 and 7dpf in *wt* and *atg13* mutant larvae, immunostained for collagen Type II (Col2a1). Scale bar = 50μm. *Right*, schematic showing where 3 measurements taken within lower jaw of larvae. (B-D) Quantification of three measurements; **(B)** length of Merkel’s cartilage, **(C)** length of lower jaw, **(D)** width of Merkel’s cartilage. n = 7 for 3dpf, n = 6 for 5dpf and n = 6 and 4 at 7dpf for *wt* and *atg13* mutant, respectively. Student’s unpaired t test performed for each age, ^ns^ *P*>.05.

Using confocal images of larval jaws labelled for Col2a1 at 3, 5 and 7dpf, the number and volume of all cells from within the cartilage elements forming the joint site was quantified (Figure 5A-C). This was achieved by utilising a modular image analysis program which can identify and outline individual Col2a1-positive cells that form the lower jaw joint during development (Figure 5B, *left*). Analysis of the collated data showed that at 5dpf and 7dpf, the number of Col2a1-positive chondrocytes forming the cartilage elements was decreased in the *atg13* mutant compared to *wt* when normalised to total element volume (Figure 5B). However, cell volume remained unchanged (Figure 5C). This could indicate decreased cell proliferation as cell volume is unaffected. To investigate this, we performed an EdU proliferation assay on larvae from 5-6dpf (Figure 5D). By analysing the number of EdU-positive chondrocytes at the joint site, we found that the *atg13* mutants showed a decrease in the number of proliferating cells within the joint compared to *wt* (Figure 5E). This implies a role for autophagy in chondrocyte development and maturation. This is of particular significance at joint sites where pre-chondrogenic cells are beginning to differentiate and mature as they are pushed up the element into the intercalation zone. Therefore, changes to differentiation and maturation here will have the biggest impact upon joint morphology.

**Figure 5.**
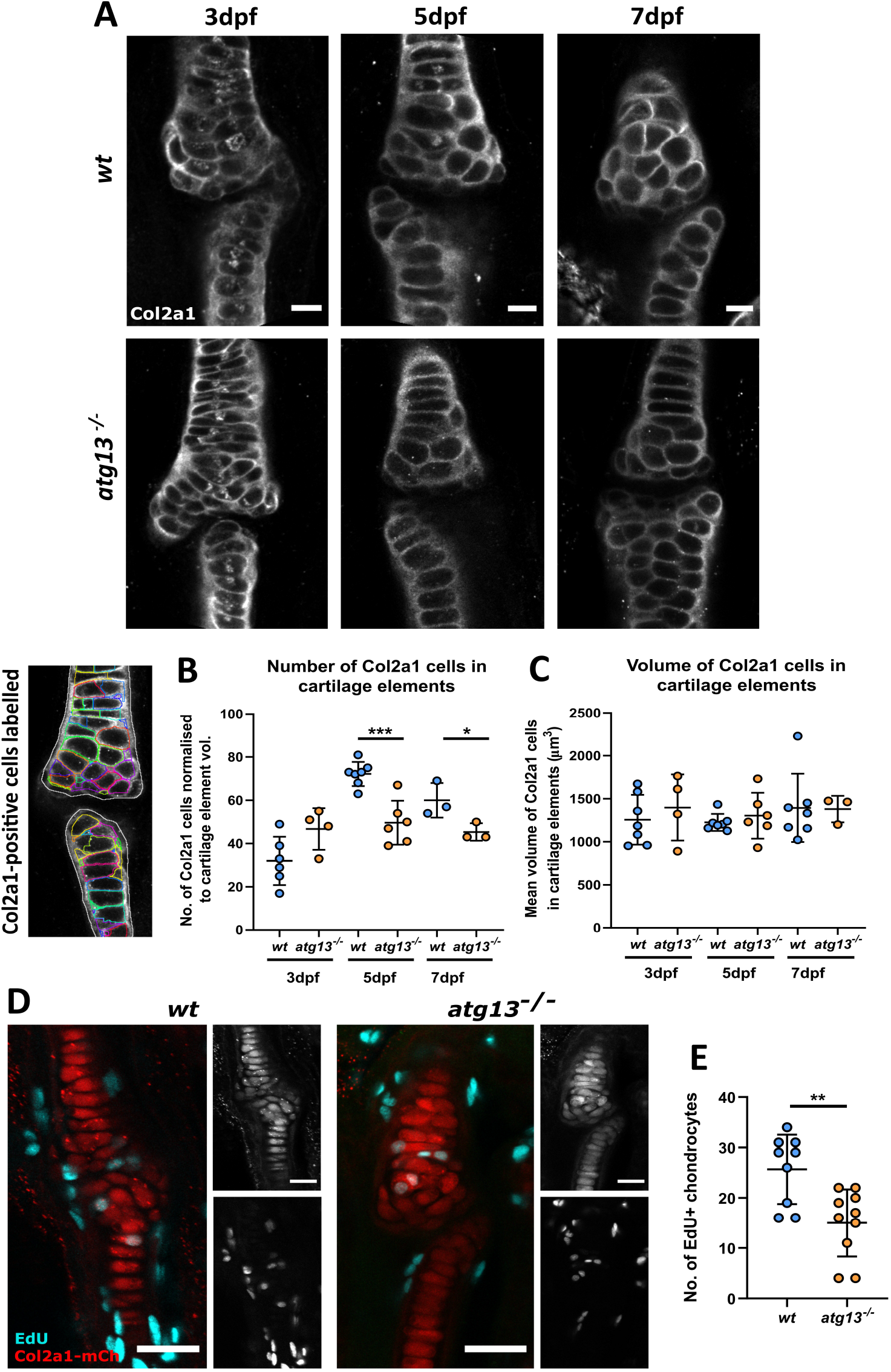
*atg13* mutant fish show decreased number of chondrocytes and reduced proliferation at joint site, **(A)** Representative confocal slices of lower jaw joint at 3, 5 and 7dpf in *wt* and *atg13* mutant fish, immunohistochemically labelled for Col2a1. Scale bars = 20μm. **(B)***Left*, example slice from confocal image showing Col2a1-positive cells outlined by modular image analysis program run in Fiji. *Right*, quantification of Col2a1-positive cell number normalised to cartilage element volume, **(C)** and volume of Col2a1-positive cells within cartilage element for *wt* and *atg13* mutants. Each data point = one larvae. Student’s unpaired t test performed where *** *P*=.0004, * *P*=.0463. **(D)** Confocal max projections of larval jaw joint in *Tg(atg13;Col2a1aBAC:mcherry) wt* and *atg13* mutants at 6dpf following 24 hour treatment with EdU Click-iT, EdU (cyan) and mCh-Col2a1 (red). Scale bars = 25μm and 20μm for insets. **(E)** Quantification of number of EdU positive chondrocytes within jaw joint region (determined as region at 5x zoom on 20x objective, when joint in middle of image plane). EdU positive chondrocytes colocalised to Col2a1 staining and counted by going through z-stack. Student’s unpaired t test performed, ** *P*=.0032.

### Chondrocytes show premature hypertrophication within a*tg13* mutants

SOX9 is one of the first transcription factors expressed within the chondrocyte maturation pathway and is essential for chondrocyte differentiation and subsequent cartilage formation ^73^. The *atg13* mutants show a significant decrease in the expression of Sox9a, an ortholog of tetrapod SOX9 ^74^, at three regions within the lower jaw: at the joint site, within the intercalation zone of the Meckel’s cartilage, and at the Meckel’s cartilage symphysis (Figure 6A-D). Thus, indicating an overall reduction in Sox9a across chondrocytes forming the lower jaw. As chondrocytes mature and become hypertrophic, Sox9a expression decreases, therefore, a reduction in Sox9a expression comparative to *wt* is indicative of premature progression of immature chondrocytes into hypertrophic chondrocytes. To confirm this, we assessed the expression of type X collagen α1 (Col10a1) via immunostaining (Figure 6E). Col10a1 can be used as a marker for hypertrophic chondrocytes, as well as early osteoblasts, as during cartilage maturation chondrocytes switch collagen expression from Col2a1 to Col10a1 ^49^. At 7dpf, we found that the *atg13* mutants had an elevated Col10a1 positive chondrocyte population compared to *wt*, indicating that these cells are maturing more rapidly. Along with reduced Sox9a data, these data indicate that loss of autophagy activity disrupts the rate of chondrocyte maturation and causes chondrocytes to become prematurely hypertrophic.

**Figure 6.**
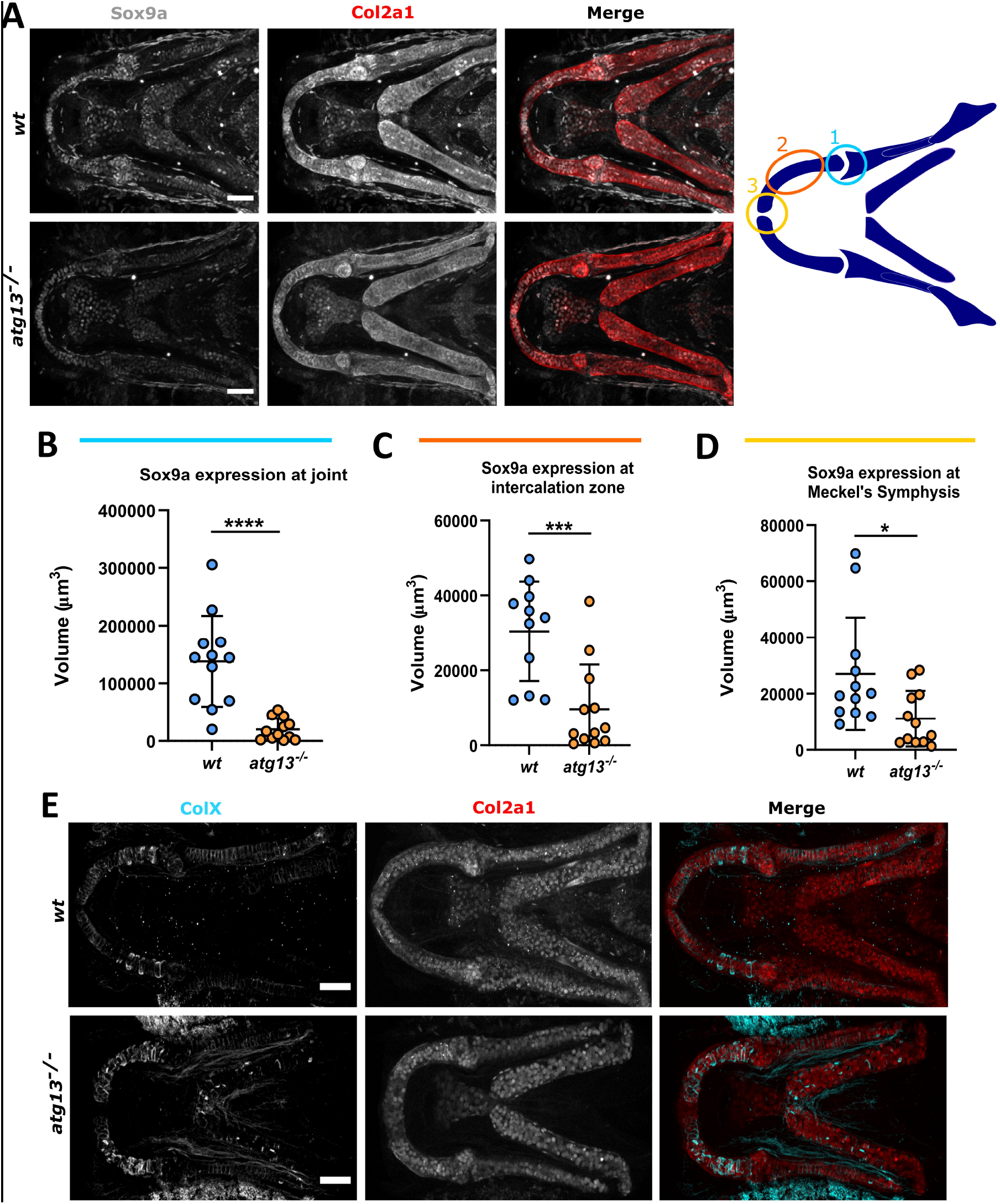
*atg13* mutation affects expression and production of key factors in cartilage development, **(A)** *Left*, confocal max projections of lower jaw at 5dpf in *wt* and *atg13* mutant fish, immunostained for Sox9a (grey) and Col2a1 (red). Scale bars = 50μm. *Right, s*chematic showing regions of interest selected within lower jaw in modular image analysis program (SoxQuant). Colours correspond to graphs below. **(B-D)** Quantification of Sox9a expression measured as volume of Sox9a within Col2a1 positive cells from confocal z-stack. Student’s unpaired t test performed where **** *P*<.0001, *** *P*=.0007, * *P*=.0173; n = 12 for both. **(E)** Confocal max projections of the lower jaw at 7dpf in *Tg(atg13;Col2a1aBAC:mcherry) wt* and *atg13* mutant larvae, immunostained for collagen Type X (ColX) (cyan) and mCherry (for Col2a1, red). Scale bars = 50μm.

### *atg13* mutant chondrocytes show alterations to chondrocyte organisation and ECM formation

To explore how these changes to chondrocyte maturation manifest within the developing tissue, we performed ultrastructural analysis on chondrocytes of the ethmoid plate at 5dpf (Figure 7). Under basal conditions, *wt* chondrocytes appeared elongated, were arranged in a stacked formation along the length of the cartilage and were surrounded by a dense and organised ECM (Figure 7A). In contrast, the *atg13* mutant chondrocytes were more disorganised with increased numbers of immature and non- or partially intercalated chondrocytes observed at the cartilage edge, resulting in a non-uniform and bumpy appearance along on the cartilage border (Figure 7B, 7D *(orange arrowheads)* and 7G). Compared to *wt*, the *atg13* mutant chondrocytes were also less electron dense and larger in size (Figure 7D), indicative of late stage hypertrophication. These data, together with the changes to sox9a and Col10a1 expression, indicate that the *atg13* mutant chondrocytes undergo an altered maturation process which inhibits proper cell placement and intercalation, leading to disorganisation of cartilage structure.

**Figure 7.**
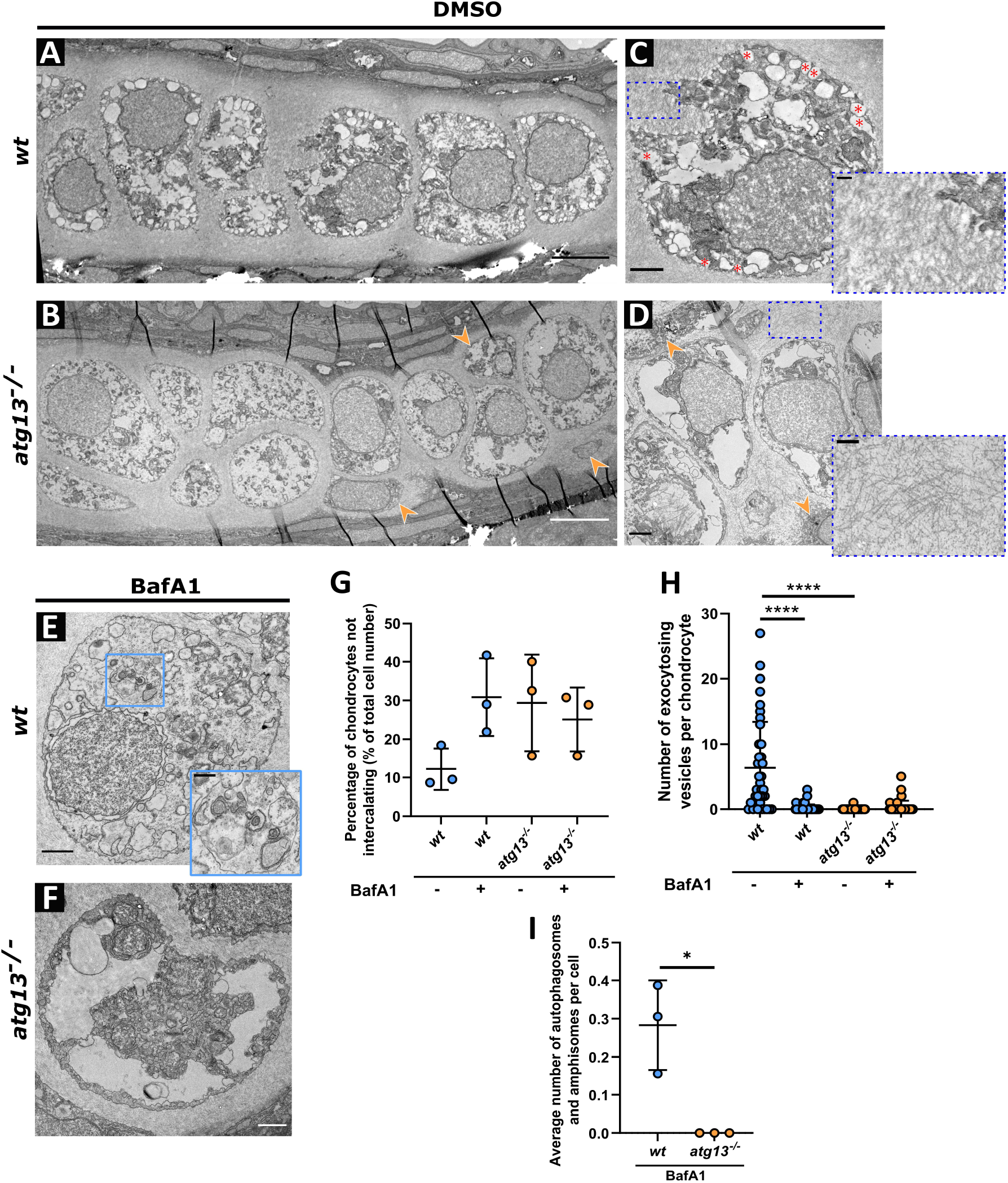
Ultrastructure and organisation of chondrocytes affected in *atg13* mutants, **(A-D)** Electron microscopy of ethmoid plate in *wt* and *atg13* mutant fish at 5dpf following DMSO or **(E, F)** BafA1 treatment for 3hrs. **(B, D)** Orange arrow heads highlight areas of non-uniformity and non-intercalating chondrocytes in *atg13* mutant cartilage. **(C)** Red asterisks show vesicles fusing with outer membrane in *wt*, not present in *atg13* mutant. Blue dotted boxes and inset show differences in ECM organisation and density between *wt* and *atg13* mutants. **(E, F)** BafA1 treatment increases number of vesicles in both *wt* and *atg13* mutants and ablates vesicle-membrane fusion events. Blue box and inset in **(E)** shows autophagosome in BafA1 treated *wt* chondrocyte, not present in *atg13* mutants. Scale bars A, B = 5μm; C, D, E, F = 1.5μm; C’, D’, E’ = 0.5μm. **(G)** Number of chondrocytes on periphery of cartilage and not aligning down central line of stack. Calculated as percentage of total cell number along ethmoid plate in one section. N = 38 chondrocytes total from 3 larvae, per condition and genotype. **(H)** Number of vesicles fusing with outer cell membrane quantified per cell following DMSO or BafA1 treatment. 2-way ANOVA performed for each; **** *P*<.0001. **(I)** Average number of autophagosomal structures per chondrocyte in bafilomycin A1 treated fish, calculated as average of all chondrocytes per individual fish. Student’s unpaired t-test performed; * *P*=.0138.

Focusing on the intracellular contents of the chondrocytes, in *wt* fish, vesicles containing low-electron dense material, predicted to be type II collagen fibrils were evident within the cytoplasm, with a substantial population fusing with the plasma membrane during exocytic events (Figure 7C, *red asterisks*). Although similar vesicles could be observed in the *atg13* mutants (Figure 7B and D), very few of those vesicles were found to be undergoing fusion with the plasma membrane (Figure 7H). This suggests that loss of autophagy affects vesicle exocytosis in chondrocytes. Notably, the ECM surrounding the chondrocytes appeared sparser and less well organised in the *atg13* mutants (Figure 7C and D, *inset*), which we hypothesise is related to reduced vesicle exocytosis, as these vesicles are predicted to contain collagens required for ECM formation.

To explore whether these effects were specific to loss of autophagy caused by the *atg13* mutation, larvae were treated with BafA1 for 3 hours at 5dpf (Figure 7E and F). We found that BafA1 treatment abrogated vesicle exocytosis in *wt* fish (Figure 7H), phenocopying the effect of the *atg13* mutation, and causing an increase in the number of vesicles within the cytoplasm of *wt* chondrocytes (Figure 7E). This supports the hypothesis that blocking or stalling of the autophagy pathway has a role in vesicle exocytosis within chondrocytes. The *wt* treated larvae also showed an increase in the number of chondrocytes not intercalating (Figure 7G), with more cells present at the outer edges of the cartilage, perpendicular to the main stack, as also seen in the mutants. These data demonstrate a role for autophagy in the control of chondrocyte differentiation and in ECM formation.

## DISCUSSION

In this study we have explored the role of autophagy in chondrocyte development and maturation. We have shown that loss of a key autophagy protein, Atg13, affects cartilage formation, joint function and is detrimental to zebrafish larval survival.

Within our model we have found that loss of atg13 affects early larval development, causing reduced growth and loss of swim bladder inflation, and juvenile lethality by 17dpf. These data demonstrate that expression of *atg13* is essential for zebrafish survival, and that Atg13 may play both autophagic and non-autophagic roles in development. This is in line with data from a previous study that showed that loss of atg13 in zebrafish affects swim bladder inflation and causes larval lethality ^25^. In a murine *Atg13^−/−^* model, Kaizuka *et al*. demonstrated that mutant mice die by embryonic day 17.5 (E17.5) and show growth retardation, as well as myocardial defects ^20^. We see no obvious changes to cardiac development within our zebrafish model and hypothesise that their delayed growth and eventual lethality is in part due to reduced yolk sac metabolism from 1-5dpf and reduced free feeding beyond 5dpf due to altered jaw morphology and function. Similarly, Mawed *et al*. demonstrated that following yolk absorption at 5dpf, *beclin1* and *atg7* knockout zebrafish mutants were unable to cope with metabolic stress and died at 9dpf and 15dpf, respectively ^63^. Defects in hepatic glycogen and lipid metabolism, and within intestinal architecture were also observed in both mutants, indicating that autophagy is critical for energy metabolism during early zebrafish development.

Through its interaction with ULK1 and FIP200, ATG13 is a vital element of the ULK1 protein kinase complex which is a key signalling node and the first protein complex within the autophagy pathway and is essential for initiating autophagosome formation ^75, 76^. Our data show that under basal conditions, the *atg13* mutant has limited autophagy activity, as indicated by a reduction in GFP-Lc3 puncta compared to *wt*, and an accumulation of p62. As an adaptor protein, p62 helps deliver cargo to the autophagosome by binding to ubiquitinated substrates and LC3, and is mainly degraded by autophagy ^77^. Therefore, an accumulation of p62 indicates autophagy inhibition. The presence of GFP-Lc3 puncta, although surprising, has been observed in other *atg13* null mouse and *C. elegans* models, and therefore, we predict could be due to LC3 aggregation or activation of a non-canonical autophagy pathway, such as LAP (Lc3-associated phagocytosis), where LC3 lipidation can occur independently of the ULK1 preinitiation complex ^78, 79^. Following treatment with Bafilomycin, the *atg13* mutants showed impaired autophagic flux as demonstrated by the minimal increase in GFP-Lc3 puncta and p62 protein levels. Using electron microscopy, we were also able to detect autophagosomes and amphisomes present in BafA1 treated *wt* chondrocytes which were completely absent in *atg13* mutant cells, further demonstrating a loss of autophagic activity caused by the *atg13* mutation (Figure 7E, *blue box inset* and 7I).

As mentioned, previous studies using mice have identified roles for autophagy in cartilage development and growth; however, the effect of these changes on joint formation and function has not been discussed. Here we have shown that *atg13* mutant zebrafish have reduced jaw function at 5dpf, which is not caused by alterations to jaw muscle development or gross jaw morphology, but instead correlates with changes to chondrocyte number at the jaw and reduced proliferation of joint precursors. These results are consistent with those from other autophagy-null murine models which also show decreased chondrocyte proliferation during development, leading to a reduction in long bone length and overall body size from birth to adulthood ^30, 50, 53^. The *atg13* mutants also showed decreased expression of an early chondrocyte marker, Sox9a. Taken together with the increased expression of Col10a1 in *atg13* mutants, these data suggest that autophagy, and specifically atg13, has a role in regulating the rate of chondrocyte maturation.

This disruption to chondrocyte maturation was confirmed by ultrastructural analysis as chondrocytes in the *atg13* mutants were more disorganised and had a reduced cellular density. Additionally, in the *atg13* mutants we observed an increase in the number of chondrocytes failing to fully intercalate into the cartilage element; a phenotype which could be induced in *wt* following treatment with BafA1, indicating that these changes may be specifically due to loss of autophagy activity. Sox9 is a key factor in early chondrogenesis and chondrocyte differentiation ^80^, and heterozygous deletion of *Sox9* in mice has been shown to cause premature chondrocyte hypertrophy and accelerated ECM mineralisation ^81^, whilst *sox9* null zebrafish show reduced chondrocyte numbers and absence of proper chondrocyte intercalation ^82^. Meanwhile, in undifferentiated mesenchymal cells, *Sox9* has been shown to be regulated by the serine/threonine protein kinase mTORC1, which is also a master regulator of autophagy ^83^. Therefore, we hypothesise that the effects on chondrocyte maturation and intercalation seen in the *atg13* mutants are due to dysregulation of *sox9a* expression, and that this dysregulation could be mediated by loss of autophagy activity.

Ultrastructural analysis revealed a decrease in ECM density and organisation in the *atg13* mutants. This is similar to the ECM phenotype seen in mice with conditional loss of *Atg7* in chondrocytes, which is caused by retention of procollagen 2 within the ER, as demonstrated by the enlarged and highly electron dense ER cisternae in the *Atg7* mutant mice ^52^. In our *atg13* mutant model, we see no obvious changes to ER structure and distribution, but we do observe a drastic reduction in the number of vesicular exocytosis events. Therefore, we hypothesise that the changes to ECM formation are due to a reduction in the secretion of collagens required for ECM formation via exocytosis. This decrease in collagen secretion could be due to their accelerated maturation, as chondrocytes reduce collagen production as they become more hypertrophic ^84, 85^. Alongside this, *Sox9* expression is required for expression of *Col2a1*, along with other collagens ^86, 87^, and therefore, its decreased expression in the *atg13* mutant could affect collagen production, leading to reduced secretion and a sparser ECM.

Here, we show that Atg13 has a role in cell differentiation during skeletal development and that changes to autophagy activity have an impact upon how joints are formed and maintained, and function. Given the link between autophagy and OA, our data highlight three possible mechanisms for increased OA risk following loss of atg13 and dysregulation of autophagy. Firstly, alterations to chondrocyte maturation can lead to altered joint function which can lead to alterations to joint loading throughout life. Secondly, premature maturation of chondrocytes could lead to increased hypertrophy of articular cartilage, leading to mineralisation, and thirdly, reduced cartilage matrix secretion could lead to cartilage that is less able to withstand physiological load and is at higher risk of breakdown. Therefore, our results identify potential links between specific autophagy proteins and cartilage health which can be used to improve our understanding of joint diseases such as OA.

## ACKNOWLEDGMENTS

We would like to acknowledge Dr Sally Hobson and Dr Chris Neal for their support in processing and imaging the EM data and the staff of the Wolfson Bioimaging Centre for confocal microscope (Leica SP5-II) access and imaging support. We would especially like to acknowledge Dr Stephen Cross for his help with image analysis and in the development of a modular analysis program. We would also like to thank Mathew Green and technical staff at the zebrafish aquarium within the University of Bristol’s Animal Scientific Unit, as well as the Aquatics BRF staff at the Francis Crick Institute for providing animal husbandry and management.

## FUNDING

J.J. Moss was funded by the Wellcome Trust Dynamic Molecular Cell Biology PhD Programme at the University of Bristol (083474). M Wirth and S.A. Tooze were supported by The Francis Crick Institute which receives its core funding from Cancer Research UK *(FC001187, FC001999)*, the UK Medical Research Council *(FC001187, FC001999)* and the Wellcome Trust *(FC001187, FC001999)*. M Wirth had been supported by a European Union Marie Curie fellowship IEF-330396. C.L. Hammond was funded by Versus Arthritis senior fellowship 21937.

## CONFLICT OF INTEREST STATEMENT

The authors declare no conflicts of interest with this article.

## AUTHOR CONTRIBUTIONS

J.J. Moss, M Wirth, S.A. Tooze, J.D. Lane and C.L. Hammond conceptualised the study and designed experiments. J.J. Moss and M Wirth conducted the experiments and interpreted data. J.J. Moss performed statistical analysis and made all figures. J.J. Moss, J.D. Lane and C.L. Hammond wrote the first draft of the manuscript and all authors made intellectual contributions and assisted in manuscript editing.

## Notes

### Competing Interest Statement

The authors have declared no competing interest.

## REFERENCES

1. Dikic I, Elazar Z. Mechanism and medical implications of mammalian autophagy. Nat Rev Mol Cell Biol. Jun 2018;19(6):349–364. doi:10.1038/s41580-018-0003-4

2. Nollet M, Santucci-Darmanin S, Breuil V, et al. Autophagy in osteoblasts is involved in mineralization and bone homeostasis. Autophagy. 2014;10(11):1965–77. doi:10.4161/auto.36182

3. Mercer TJ, Gubas A, Tooze SA. A molecular perspective of mammalian autophagosome biogenesis. Journal of biological chemistry. 2018;293(15):5386–5395.

4. Mizushima N. The role of the Atg1/ULK1 complex in autophagy regulation. Current opinion in cell biology. 2010;22(2):132–139.

5. Chan EY, Kir S, Tooze SA. siRNA screening of the kinome identifies ULK1 as a multidomain modulator of autophagy. Journal of Biological Chemistry. 2007;282(35):25464–25474.

6. Ganley IG, Lam DH, Wang J, Ding X, Chen S, Jiang X. ULK1· ATG13· FIP200 complex mediates mTOR signaling and is essential for autophagy. Journal of Biological Chemistry. 2009;284(18):12297–12305.

7. Hieke N, Loffler AS, Kaizuka T, et al. Expression of a ULK1/2 binding-deficient ATG13 variant can partially restore autophagic activity in ATG13-deficient cells. Autophagy. 2015;11(9):1471–83. doi:10.1080/15548627.2015.1068488

8. Qi S, Stjepanovic G, Hurley JH. Structure of the human Atg13-Atg101 HORMA heterodimer: an interaction hub within the ULK1 complex. Structure. 2015;23(10):1848–1857.

9. Hegedűs K, Nagy P, Gáspári Z, Juhász G. The putative HORMA domain protein Atg101 dimerizes and is required for starvation-induced and selective autophagy in Drosophila. BioMed research international. 2014;2014

10. Jao CC, Ragusa MJ, Stanley RE, Hurley JH. A HORMA domain in Atg13 mediates PI 3-kinase recruitment in autophagy. Proceedings of the National Academy of Sciences. 2013;110(14):5486–5491.

11. Park J-M, Jung CH, Seo M, et al. The ULK1 complex mediates MTORC1 signaling to the autophagy initiation machinery via binding and phosphorylating ATG14. Autophagy. 2016;12(3):547–564.

12. Chang Y-Y, Neufeld TP. An Atg1/Atg13 complex with multiple roles in TOR-mediated autophagy regulation. Molecular biology of the cell. 2009;20(7):2004–2014.

13. Tian E, Wang F, Han J, Zhang H. epg-1 functions in autophagy-regulated processes and may encode a highly divergent Atg13 homolog in C. elegans. Autophagy. 2009;5(5):608–615.

14. Kraft C, Kijanska M, Kalie E, et al. Binding of the Atg1/ULK1 kinase to the ubiquitin□like protein Atg8 regulates autophagy. The EMBO journal. 2012;31(18):3691–3703.

15. Karanasios E, Stapleton E, Manifava M, et al. Dynamic association of the ULK1 complex with omegasomes during autophagy induction. Journal of cell science. 2013;126(22):5224–5238.

16. Hosokawa N, Hara T, Kaizuka T, et al. Nutrient-dependent mTORC1 association with the ULK1–Atg13–FIP200 complex required for autophagy. Molecular biology of the cell. 2009;20(7):1981–1991.

17. Alers S, Löffler AS, Paasch F, et al. Atg13 and FIP200 act independently of Ulk1 and Ulk2 in autophagy induction. Autophagy. 2011;7(12):1424–1433.

18. Alers S, Wesselborg S, Stork B. ATG13: just a companion, or an executor of the autophagic program? Autophagy. Jun 2014;10(6):944–56. doi:10.4161/auto.28987

19. Gan B, Peng X, Nagy T, Alcaraz A, Gu H, Guan J-L. Role of FIP200 in cardiac and liver development and its regulation of TNFα and TSC–mTOR signaling pathways. The Journal of cell biology. 2006;175(1):121–133.

20. Kaizuka T, Mizushima N. Atg13 is essential for autophagy and cardiac development in mice. Molecular and cellular biology. 2016;36(4):585–595.

21. Komatsu M, Waguri S, Ueno T, et al. Impairment of starvation-induced and constitutive autophagy in Atg7-deficient mice. Journal of Cell Biology. 2005;169(3):425–434.

22. Yue Z, Jin S, Yang C, Levine AJ, Heintz N. Beclin 1, an autophagy gene essential for early embryonic development, is a haploinsufficient tumor suppressor. Proceedings of the National Academy of Sciences. 2003;100(25):15077–15082.

23. Zhou X, Takatoh J, Wang F. The mammalian class 3 PI3K (PIK3C3) is required for early embryogenesis and cell proliferation. PloS one. 2011;6(1):e16358.

24. Kuma A, Komatsu M, Mizushima N. Autophagy-monitoring and autophagy-deficient mice. Autophagy. 2017;13(10):1619–1628. doi:10.1080/15548627.2017.1343770

25. Morishita H, Kanda Y, Kaizuka T, et al. Autophagy Is Required for Maturation of Surfactant-Containing Lamellar Bodies in the Lung and Swim Bladder. Cell reports. 2020;33(10):108477.

26. Cadwell K, Debnath J. Beyond self-eating: The control of nonautophagic functions and signaling pathways by autophagy-related proteins. Journal of Cell Biology. 2018;217(3):813–822.

27. Neufeld TP. Autophagy and cell growth–the yin and yang of nutrient responses. Journal of cell science. 2012;125(10):2359–2368.

28. Deegan S, Saveljeva S, Gorman AM, Samali A. Stress-induced self-cannibalism: on the regulation of autophagy by endoplasmic reticulum stress. Cellular and Molecular Life Sciences. 2013;70(14):2425–2441.

29. Mizushima N, Levine B. Autophagy in mammalian development and differentiation. Nature cell biology. 2010;12(9):823–830.

30. Horigome Y, Ida-Yonemochi H, Waguri S, Shibata S, Endo N, Komatsu M. Loss of autophagy in chondrocytes causes severe growth retardation. Autophagy. Mar 2020;16(3):501–511. doi:10.1080/15548627.2019.1628541

31. Chang J, Wang W, Zhang H, Hu Y, Wang M, Yin Z. The dual role of autophagy in chondrocyte responses in the pathogenesis of articular cartilage degeneration in osteoarthritis. Int J Mol Med. Dec 2013;32(6):1311–8. doi:10.3892/ijmm.2013.1520

32. Carames B, Taniguchi N, Otsuki S, Blanco FJ, Lotz M. Autophagy is a protective mechanism in normal cartilage, and its aging-related loss is linked with cell death and osteoarthritis. Arthritis Rheum. Mar 2010;62(3):791–801. doi:10.1002/art.27305

33. Levine B, Kroemer G. Biological functions of autophagy genes: a disease perspective. Cell. 2019;176(1-2):11–42.

34. Bouderlique T, Vuppalapati KK, Newton PT, Li L, Barenius B, Chagin AS. Targeted deletion of Atg5 in chondrocytes promotes age-related osteoarthritis. Annals of the rheumatic diseases. 2016;75(3):627–631.

35. Jeon H, Im GI. Autophagy in osteoarthritis. Connect Tissue Res. Nov 2017;58(6):497–508. doi:10.1080/03008207.2016.1240790

36. Duan R, Xie H, Liu Z-Z. The Role of Autophagy in Osteoarthritis. Frontiers in Cell and Developmental Biology. 2020;8:1437.

37. Murray CJ, Lopez AD. Alternative projections of mortality and disability by cause 1990–2020: Global Burden of Disease Study. The lancet. 1997;349(9064):1498–1504.

38. Woo T, Lau L, Cheung N, Chan P, Tan K, Gardner A. Efficacy of oral collagen in joint pain—osteoarthritis and rheumatoid arthritis. J Arthritis. 2017;6(233):2.

39. Neogi T. The epidemiology and impact of pain in osteoarthritis. Osteoarthritis and cartilage. 2013;21(9):1145–1153.

40. Li YS, Zhang FJ, Zeng C, et al. Autophagy in osteoarthritis. Joint Bone Spine. Mar 2016;83(2):143–8. doi:10.1016/j.jbspin.2015.06.009

41. Litwic A, Edwards MH, Dennison EM, Cooper C. Epidemiology and burden of osteoarthritis. British medical bulletin. 2013;105(1):185–199.

42. Faber BG, Baird D, Gregson CL, et al. DXA-derived hip shape is related to osteoarthritis: findings from in the MrOS cohort. Osteoarthritis and cartilage. 2017;25(12):2031–2038.

43. Waarsing J, Rozendaal R, Verhaar J, Bierma-Zeinstra S, Weinans H. A statistical model of shape and density of the proximal femur in relation to radiological and clinical OA of the hip. Osteoarthritis and cartilage. 2010;18(6):787–794.

44. Beck M, Kalhor M, Leunig M, Ganz R. Hip morphology influences the pattern of damage to the acetabular cartilage: femoroacetabular impingement as a cause of early osteoarthritis of the hip. The Journal of bone and joint surgery British volume. 2005;87(7):1012–1018.

45. Baker-LePain JC, Lane NE. Relationship between joint shape and the development of osteoarthritis. Current opinion in rheumatology. 2010;22(5):538.

46. Provot S, Schipani E. Molecular mechanisms of endochondral bone development. Biochemical and biophysical research communications. 2005;328(3):658–665.

47. Dodds G. Row formation and other types of arrangement of cartilage cells in endochondral ossification. The Anatomical Record. 1930;46(4):385–399.

48. Davidson LA, Joshi SD, Kim HY, Von Dassow M, Zhang L, Zhou J. Emergent morphogenesis: elastic mechanics of a self-deforming tissue. Journal of biomechanics. 2010;43(1):63–70.

49. Akkiraju H, Nohe A. Role of chondrocytes in cartilage formation, progression of osteoarthritis and cartilage regeneration. Journal of developmental biology. 2015;3(4):177–192.

50. Vuppalapati KK, Bouderlique T, Newton PT, et al. Targeted deletion of autophagy genes Atg5 or Atg7 in the chondrocytes promotes caspase□dependent cell death and leads to mild growth retardation. Journal of Bone and Mineral Research. 2015;30(12):2249–2261.

51. Srinivas V, Bohensky J, Shapiro IM. Autophagy: a new phase in the maturation of growth plate chondrocytes is regulated by HIF, mTOR and AMP kinase. Cells Tissues Organs. 2009;189(1-4):88–92.

52. Cinque L, Forrester A, Bartolomeo R, et al. FGF signalling regulates bone growth through autophagy. Nature. 2015;528(7581):272–275.

53. Kang X, Yang W, Feng D, et al. Cartilage□specific autophagy deficiency promotes ER stress and impairs chondrogenesis in PERK□ATF4□CHOP–dependent manner. Journal of Bone and Mineral Research. 2017;32(10):2128–2141.

54. Aleström P, D’Angelo L, Midtlyng PJ, et al. Zebrafish: Housing and husbandry recommendations. Laboratory Animals. 2020;54(3):213–224.

55. He C, Bartholomew CR, Zhou W, Klionsky DJ. Assaying autophagic activity in transgenic GFP-Lc3 and GFP-Gabarap zebrafish embryos. Autophagy. 2009;5(4):520–526.

56. Hammond CL, Schulte-Merker S. Two populations of endochondral osteoblasts with differential sensitivity to Hedgehog signalling. Development. 2009;136(23):3991–4000.

57. Jao L-E, Wente SR, Chen W. Efficient multiplex biallelic zebrafish genome editing using a CRISPR nuclease system. Proceedings of the National Academy of Sciences. 2013;110(34):13904–13909.

58. Dahlem TJ, Hoshijima K, Jurynec MJ, et al. Simple methods for generating and detecting locus-specific mutations induced with TALENs in the zebrafish genome. PLoS Genet. 2012;8(8):e1002861.

59. Wilkinson RN, Elworthy S, Ingham PW, van Eeden FJ. A method for high-throughput PCR-based genotyping of larval zebrafish tail biopsies. Biotechniques. 2013;55(6):314–316.

60. Schindelin J, Arganda-Carreras I, Frise E, et al. Fiji: an open-source platform for biological-image analysis. Nature methods. 2012;9(7):676–682.

61. MIA: Version 0.9.30. 2019. https://github.com/SJCross/MIA

62. Brunt LH, Begg K, Kague E, Cross S, Hammond CL. Wnt signalling controls the response to mechanical loading during zebrafish joint development. Development. 2017;144(15):2798–2809.

63. Mawed SA, Zhang J, Ren F, He Y, Mei J. atg7 and beclin1 are essential for energy metabolism and survival during the larval-to-juvenile transition stage of zebrafish. Aquaculture and Fisheries. 2021;

64. Klionsky DJ, Abdelmohsen K, Abe A, et al. Guidelines for the use and interpretation of assays for monitoring autophagy. Autophagy. 2016;12(1):1–222.

65. Tian Y, Li Z, Hu W, et al. C. elegans screen identifies autophagy genes specific to multicellular organisms. Cell. 2010;141(6):1042–1055.

66. Liang Q, Yang P, Tian E, Han J, Zhang H. The C. elegans ATG101 homolog EPG-9 directly interacts with EPG-1/Atg13 and is essential for autophagy. Autophagy. 2012;8(10):1426–1433.

67. Chávez MN, Morales RA, López-Crisosto C, Roa JC, Allende ML, Lavandero S. Autophagy Activation in Zebrafish Heart Regeneration. Scientific reports. 2020;10(1):1–11.

68. Sasaki T, Lian S, Qi J, et al. Aberrant autolysosomal regulation is linked to the induction of embryonic senescence: differential roles of Beclin 1 and p53 in vertebrate Spns1 deficiency. PLoS Genet. Jun 2014;10(6):e1004409. doi:10.1371/journal.pgen.1004409

69. Santos-Ledo A, Garcia-Macia M, Campbell PD, Gronska M, Marlow FL. Kinesin-1 promotes chondrocyte maintenance during skeletal morphogenesis. PLoS Genetics. 2017;13(7):e1006918.

70. Schilling TF, Kimmel CB. Musculoskeletal patterning in the pharyngeal segments of the zebrafish embryo. Development. 1997;124(15):2945–2960.

71. Askary A, Smeeton J, Paul S, et al. Ancient origin of lubricated joints in bony vertebrates. Elife. 2016;5:e16415.

72. Hernández LP, Barresi MJ, Devoto SH. Functional morphology and developmental biology of zebrafish: reciprocal illumination from an unlikely couple. Integrative and comparative biology. 2002;42(2):222–231.

73. Bi W, Deng JM, Zhang Z, Behringer RR, De Crombrugghe B. Sox9 is required for cartilage formation. Nature genetics. 1999;22(1):85–89.

74. Chiang EF-L, Pai C-I, Wyatt M, Yan Y-L, Postlethwait J, Chung B-c. Two sox9 genes on duplicated zebrafish chromosomes: expression of similar transcription activators in distinct sites. Developmental biology. 2001;231(1):149–163.

75. Lane JD, Korolchuk VI, Murray JT, Zachari M, Ganley IG. The mammalian ULK1 complex and autophagy initiation. Essays in biochemistry. 2017;61(6):585–596.

76. Itakura E, Mizushima N. Characterization of autophagosome formation site by a hierarchical analysis of mammalian Atg proteins. Autophagy. 2010;6(6):764–776.

77. Pankiv S, Clausen TH, Lamark T, et al. p62/SQSTM1 binds directly to Atg8/LC3 to facilitate degradation of ubiquitinated protein aggregates by autophagy. Journal of biological chemistry. 2007;282(33):24131–24145.

78. Henault J, Martinez J, Riggs JM, et al. Noncanonical autophagy is required for type I interferon secretion in response to DNA-immune complexes. Immunity. 2012;37(6):986–997.

79. Kim J-Y, Zhao H, Martinez J, et al. Noncanonical autophagy promotes the visual cycle. Cell. 2013;154(2):365–376.

80. Kronenberg HM. Developmental regulation of the growth plate. Nature. 2003;423(6937):332–336.

81. Akiyama H, Chaboissier M-C, Martin JF, Schedl A, de Crombrugghe B. The transcription factor Sox9 has essential roles in successive steps of the chondrocyte differentiation pathway and is required for expression of Sox5 and Sox6. Genes & development. 2002;16(21):2813–2828.

82. Yan Y-L, Willoughby J, Liu D, et al. A pair of Sox: distinct and overlapping functions of zebrafish sox9 co-orthologs in craniofacial and pectoral fin development. Development. 2005;132(5):1069–1083.

83. Lane JD, Korolchuk VI, Murray JT, Rabanal-Ruiz Y, Otten EG. mTORC1 as the main gateway to autophagy. Essays in biochemistry. 2017;61(6):565–584.

84. Samsa WE, Zhou X, Zhou G. Signaling pathways regulating cartilage growth plate formation and activity. Elsevier; 2017:3–15.

85. Dreier R. Hypertrophic differentiation of chondrocytes in osteoarthritis: the developmental aspect of degenerative joint disorders. Arthritis research & therapy. 2010;12(5):1–11.

86. Lefebvre V, Behringer R, De Crombrugghe B. L-Sox5, Sox6 and Sox9 control essential steps of the chondrocyte differentiation pathway. Osteoarthritis and cartilage. 2001;9:S69–S75.

87. Ng L-J, Wheatley S, Muscat GE, et al. SOX9 binds DNA, activates transcription, and coexpresses with type II collagen during chondrogenesis in the mouse. Developmental biology. 1997;183(1):108–121.

