## Supplemental Figures S1 and S4 for "Autophagy coordinates chondrocyte development and early joint formation in zebrafish"

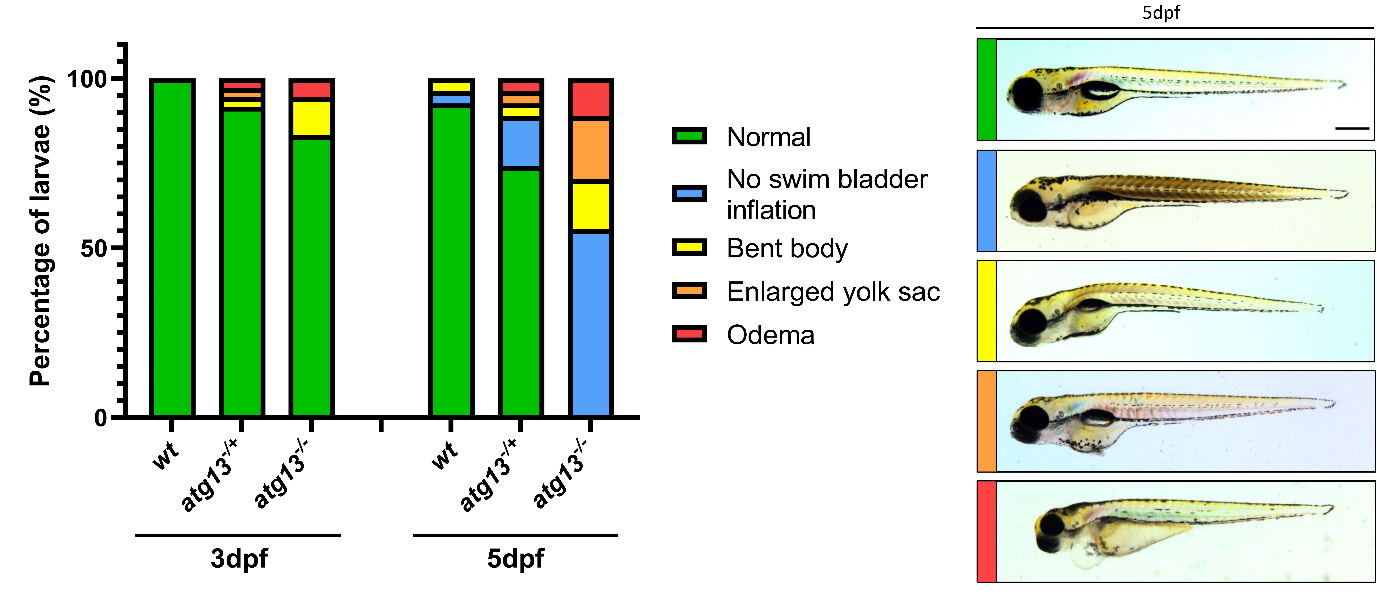

**Supplemental figures S2 and S3 – *atg13* mutants show reduced jaw function,** Video clips of larval jaw movements from 5dpf *wt* (Supplemental figure S2) and *atg13* mutant (Supplemental figure S3) larvae. Video clips taken from longer videos filmed at 1 frame per second for 1000 frames.

**Supplemental Figure S1 – Loss of *atg13* causes developmental defects from 3dpf in zebrafish larvae**, *Left,* quantification of phenotypic differences observed at 3dpf and 5dpf in *wt,* heterozygous and homozygous mutant *atg13* larvae. Larvae at 3dpf show no swim bladder inflation but this is normal at this stage of development. *Right,* representative brightfield images, taken at 5dpf, show an example of each developmental defect with severity categorised by colour (green = healthy, blue-orange = moderate, red = severe). N = 24 for both ages and all genotypes. Scale bar = 500µm.

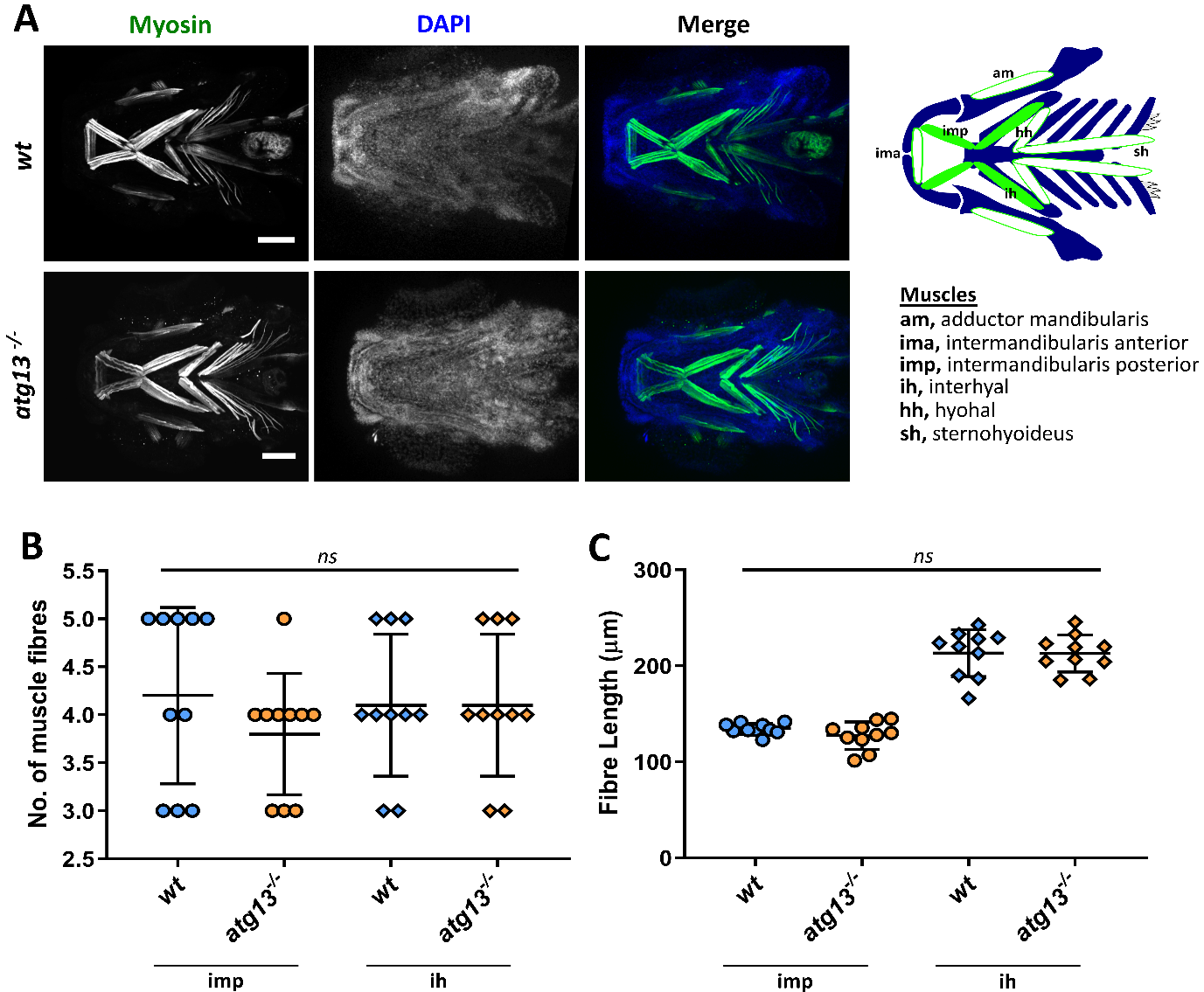

**Supplemental Figure S4 – *atg13* mutation does not affect muscle development in the lower jaw, (A)** *Left,* representative confocal z-stack projections of the lower jaw at 5dpf in *wt* and *atg13* mutant larvae, immunostained for muscle (green) and counterstained with DAPI (blue). Scale bar = 100µm. *Right,* schematic showing key muscle groups (orange) within the lower jaw at 5dpf. Quantification of muscle fibre number **(B)** and muscle length **(C)** of IMP, IH and IMA muscles of *wt* and *atg13* mutant fish, n = 5 for all, right and left side counted separately per fish. *IMP* = intermandibularis posterior, *IH* = interhyal, *IMA* = intermandibularis anterior.
